# Gene Expression and Evolution in the Smalltooth Sawfish, *Pristis pectinata*

**DOI:** 10.1101/2023.01.12.523848

**Authors:** Taiya M. Jarva, Nicole M. Phillips, Cory Von Eiff, Gregg R. Poulakis, Gavin Naylor, Kevin A. Feldheim, Alex S. Flynt

## Abstract

Sawfishes (Pristidae) are large, highly threatened rays named for their tooth-studded rostrum, which is used for prey sensing and capture. Of all five species, the smalltooth sawfish, *Pristis pectinata*, has experienced the greatest decline in range, currently found in only ∼20% of its historic range. To better understand the genetic underpinnings of these taxonomically and morphologically unique animals, we collected transcriptomic data from several tissue types, mapped them to the recently completed reference genome and contrasted the patterns observed with comparable data from other elasmobranchs. Evidence of positive selection was detected in 79 genes in *P. pectinata*, several of which are involved in growth factor/receptor tyrosine kinase signaling and specification of organ symmetry, suggesting a role in morphogenesis. Data acquired also allow for examination of the molecular components of *P. pectinata* electrosensory systems, which are highly developed in sawfishes and have likely been influential in their evolutionary success.

---

As meso-to apex-level predators, chondrichthyans (sharks, rays, and chimaeras) play essential roles in the health of marine ecosystems by increasing biodiversity, buffering against invasive species, decreasing transmission of diseases, and mitigating the effects of climate change^1,2^ However, many chondrichthyan populations are in decline, primarily due to overfishing, with more than one third estimated to be threatened with extinction^3^. Sawfishes belong to one of the most threatened families, with all five species assessed as Endangered or Critically Endangered on the International Union for the Conservation of Nature (IUCN) Red List of Threatened Species^4^. Sawfishes are notable for their tooth-studded rostrum, which is used to detect, acquire, and manipulate prey^5,6^. This rostrum also makes them especially susceptible to entanglement in fishing gear, which has precipitated a global decline in their numbers over the last century.^4,7,8^

Bycatch in fisheries and habitat degradation continue to pose the greatest threats to sawfishes, including in ‘stronghold’ locations such as the United States and Australia (NMFS 2009)^9^. Fisher education and trade bans on sawfish and their parts have been used to mitigate sawfish declines^10,8^. While such strategies are essential, additional preventative initiatives are needed as sawfish populations continue to decline globally^11^. Recent research aimed at supporting development of deterrent technology found that sawfish will react to electric field stimuli, but behavioral responses were not consistent and were considered insufficient to avoid fisheries gear entanglement^12^.

The electrosensing abilities of sawfishes are currently poorly understood, and little is known about the underlying the molecular mechanisms of electroreception and environmental biosensing that underpin their behaviors. The aims of this study were to collect transcriptome sequences from different tissues of the Critically Endangered smalltooth sawfish, *Pristis pectinata*, map the data to the recently completed high resolution genome assembly, and compare patterns of gene expression and sequence evolution with those derived from other elasmobranchs to identify unique components of genomic architecture. Of all sawfishes, *P. pectinata* has experienced the most severe decline in range and is present in less than 20% of its former range in the Atlantic Ocean^7,8^. Viable populations are currently restricted to Florida in the U.S. and western portions of The Bahamas and, like all sawfishes, reducing fisheries interactions is a top conservation priority^4^. The gene set collected in this study provides insight into evolutionary patterns and the mechanisms of electrosensing in *P. pectinata* and serve as critical first steps for future work.

## High quality *Pristis pectinata* gene set

In collaboration with the Vertebrate Genome Project, GN produced a high-quality reference genome assembly for an adult female *P. pectinata* that had died at the Ripley’s Aquarium in Myrtle Beach, South Carolina (GCA_009764475.1). The assembly was based on 60x PACBIO long read sequencing, BioNano, Hi-C and Illumina short read data (scaffold N50 101.7M; Contig N50=17M; 99.61% of the data assigned to 48 chromosomes. Genome size 2.27Gb; http://vertebrategenomesproject.org). However, while the assembly had high contiguity, there were limited EST data associated with the original genome assembly^13^. As a result, approximately 40% of expected orthologs were absent from the predicted gene dataset (Fig. 1E)^14^. To address this issue and maximize utility of the *P. pectinata* genome, RNA was sequenced from tissues collected from a juvenile female (828 mm stretch total length) collected by GP under ESA Permit No. 21043. Datasets were generated from brain, kidney, liver, ovary, and skin tissues fixed in RNAlater.

**Figure 1.**
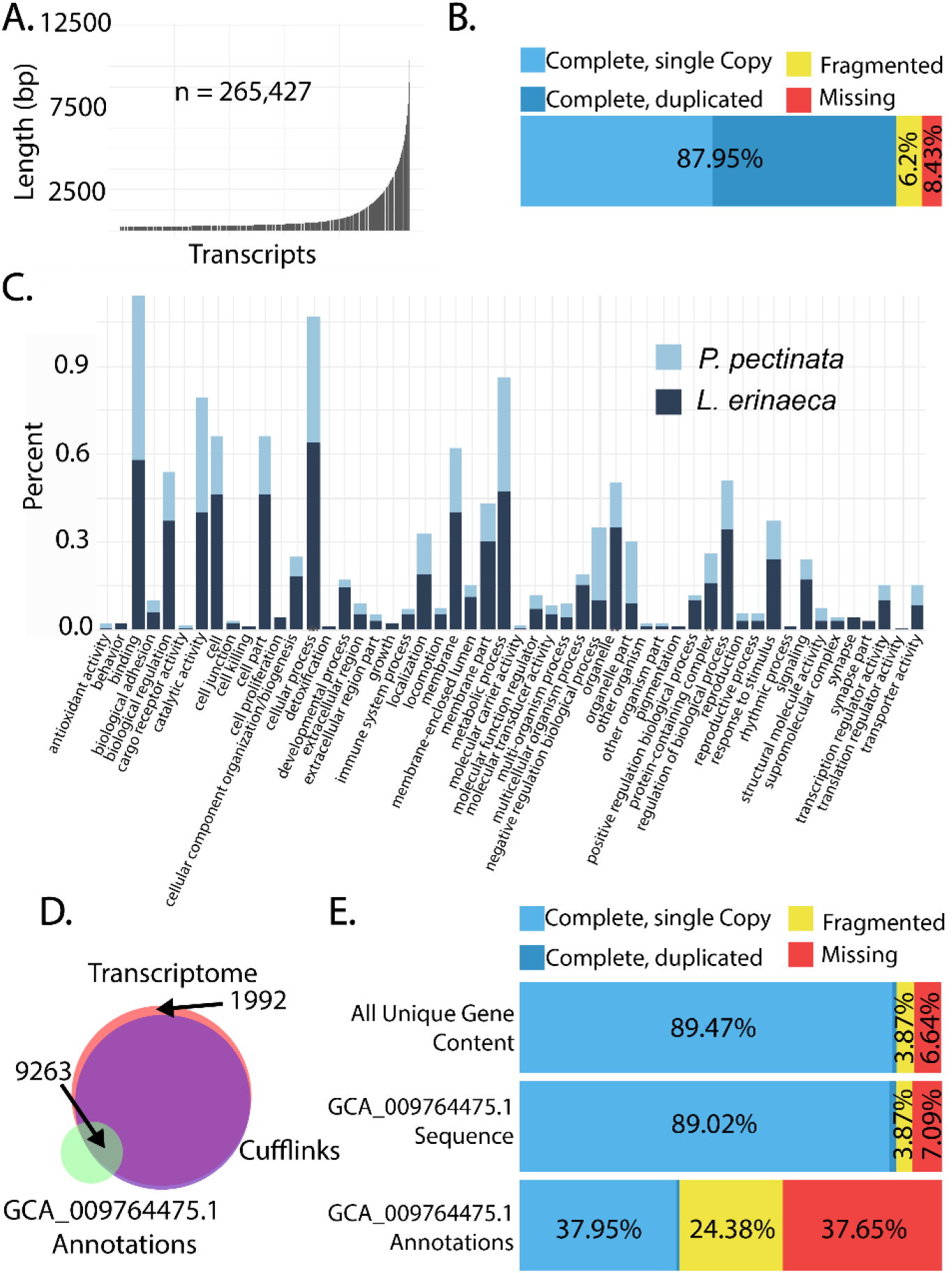
Establishment of a near complete gene set for the smalltooth sawfish, *Pristis pectinata*. **A)** Log length distribution of transcripts in assembled transcriptome with total number of transcripts (n = 265,427). **B)** Outcome of Benchmarking Universal Single Copy Ortholog (BUSCO) analysis to assess completeness of assembled transcriptome. **C)** WeGO (Ye et al., 2018) gene ontology analysis of *P. pectinata* and little skate, *Leucoraja erinacea*, transcriptomes. Genes are classified according to biological process, molecular function, or cellular component and plotted by percent of genes related to meaningful biological activities. *P. pectinata* has a higher percentage of genes related to DNA binding, negative regulation of biological processes, organelle part, and antioxidant activity, while *L. erinacea* has more genes related to cellular processes, response to stimulus, signaling, and catalytic activity. **D)** Venn diagram (Hulsen et al., 2008) showing overlap of the number of transcripts provided with the public genome (GCA_009764475.1, green), the Cufflinks assisted annotation of the public genome (purple), and transcripts in the de novo assembled transcriptome (pink). **E)** BUSCO assessment for all unique gene content from combined transcriptome and genome recourses, and annotations associated with GCA_009764475.1.

Genome annotation facilitated by RNA-seq alignments yielded substantially more transcripts (159,014), than the 19,597 genes predicted from the genome alone. RNA-seq data were also assembled into a *de novo* transcriptome of 265,427 transcripts with 31.9% being >500 bp (Fig. 1A-B). Substantial improvement was seen in the contribution of the *de novo* transcriptome and RNA-seq guided annotation, with approximately 2000 genes exclusive to the *de novo* transcriptome. Less redundancy was seen in the dataset after intersecting *de novo* transcripts with predicted genes from the genome, with 9,233 transcripts shared between datasets (Fig. 1D). Combining the reference genome assembly predictions, RNA-seq assisted annotations, and a *de novo* transcriptome resulted in a greatly enhanced gene set representation (Fig. 1E). In the *de novo* transcriptome alone, less than 10% of expected orthologs were absent. In the combined gene set, the percentage of missing genes was reduced to only 6.6% (Sup. Tbl. 2).

To further validate the *P. pectinata* gene set, gene ontology (GO) terms were assigned and compared to the transcriptome of the little skate, *Leucoraja erinacea*^15^ (Fig. 1C). Distributions of high-level GO terms were similar between *P. pectinata* and *L. erinacea*, suggesting gene content in the *de novo* transcriptome represents what is observed in related taxa. However, differences were noted, such as a higher percentage of genes involved in antioxidant activity and DNA binding in *P. pectinata* versus a higher percentage of genes in *L. erinacea* related to signaling, response to stimulus, metabolic process, and catalytic activity. This may reflect a difference in juvenile and adult tissues sampled for *P. pectinata* relative to the embryonic tissue-derived *L. erinacea* transcriptome. The near-complete *P. pectinata* gene set enables characterization of unique genetics in this species, which was not possible with predicted annotations offered by genome sequence approaches alone.

## Unique genetic features of *Pristis pectinata*

To identify positively selected genes (PSGs) in *P. pectinata*, the combined dataset from *P. pectinata* and transcriptomes of four other fish species were assigned to orthologous gene groups with Orthofinder^16^. The species included were the Australian ghostshark, *Callorhinchus milii*, chain catshark, *Scyliorhinus retifer*, and *L. erinacea*, with the Indonesian coelacanth, *Latimeria menadoensis*, serving as an outgroup (Fig. 2A). *Pristis pectinata* had the second highest percentage of genes (∼51%) assigned to an orthogroup and the highest percentage of species-specific genes, likely due to the substantially greater completeness of the assembly relative to the other species (Sup. Fig. 1). The resulting 3,116 genes were tested for branch-specific episodic selection using aBSREL, revealing 79 PSGs in *P. pectinata*^17^ (Supplementary Methods, Sup. Tbl. 3).

**Figure 2.**
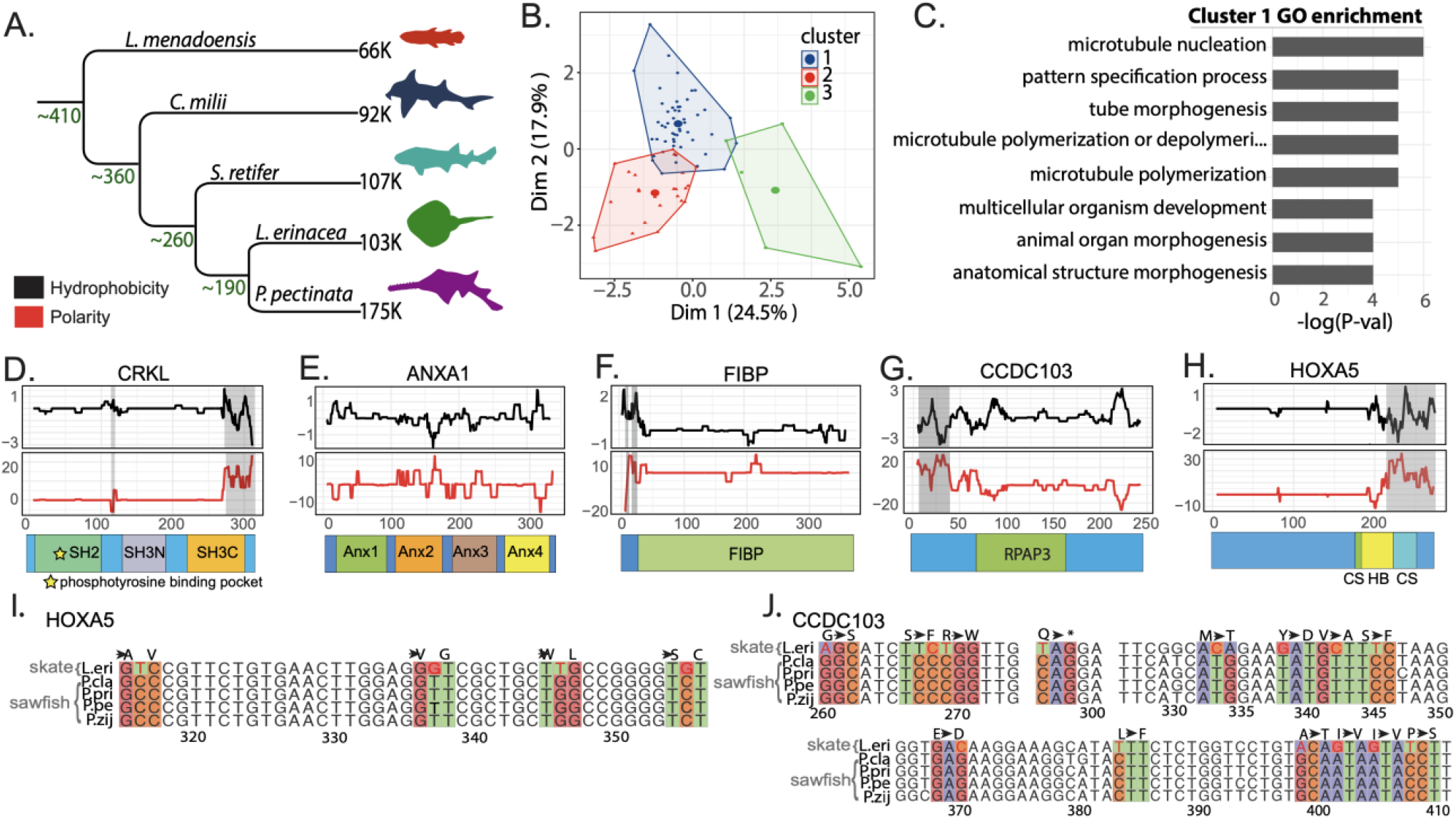
Smalltooth sawfish, *Pristis pectinata*, genes under positive selection. **A)** Phylogeny of taxa used in positive selection analysis with approximate divergence times shown at each node and number of transcripts in each transcriptome noted at each terminal branch. Species included: *Latimeria menadoensis, Callorhinchus milii, Scyliorhinus retifer, Leucoraja erinacea*, and *P. pectinata*. **B)** Results of k-means clustering analysis of selected genes using omega value, percent of sites, change in GRAVY value relative to little skate, *L. erinacea*, and change in expression relative to *S. retifer*. **C)** Top 8 most enriched gene ontology (GO) terms related to biological processes from genes which were grouped into Cluster 1 by PCA analysis plotted by -log(p-value). **D-H)** Changes in hydrophobicity and polarity of protein sequence relative to *L. erinacea*. Sequence alignment gaps are shown by shaded regions. Functional protein domains retrieved from InterproScan and literature are shown as colored boxes below each plot. **I)** Sequence alignment of PCR-amplified region of interest from HoxA5 between *Pristis* sawfishes and *L. erinacea* with codons of interest colored by nucleotide. **J)** Alignment of region of interest from amplification of CCDC103.Alignments are colored by nucleotide.

To cluster *P. pectinata* PSGs into groups which may share functional relationships or similar selection pressures, omega values from aBSREL, percent of sites under selection, gene expression rank relative to *S. retifer* orthologs, and the difference in Grand Average of Hydropathy (GRAVY) between *P. pectinata* and *L. erinacea* orthologs were collected for Principal Components Analysis (Sup. File 1). K-means clustering identified three groups containing 49, 25, and 5 genes (Fig. 2B). Generally, changes in GRAVY and omega values were highly correlated, with GRAVY values being the major contribution to variance (Sup. Fig. 2A). Between tissues, the highest correlation was seen between skin, kidney, and brain, while liver and ovary showed little correlation.

Gene ontology enrichment analysis by TopGO of PSGs in Cluster 1 versus all genes analyzed showed enrichment in functions such as multicellular organism development, animal organ morphogenesis, and anatomical structure morphogenesis^18^ (Fig. 2C). In comparison, Cluster 2 genes were related to regulation of response to biotic stimulus, regulation of defense response, and proteasomal protein catabolic process (Sup. Fig. 2B). Interestingly, Cluster 1 contained multiple genes implicated in developmental processes such as fibroblast growth factor/receptor tyrosine kinase (FGF/RTK) and mitogen activated protein kinase (MAPK) signaling, suggesting that *P. pectinata* PSGs share functional or evolutionary similarities. The altered FGF and MAPK signaling that may result from these diverged genes implies possible relevance to *P. pectinata*-specific morphogenesis.

Comparing biochemical properties such as hydrophobicity and polarity of developmental genes with the closest available taxon, *L. erinacea*, revealed functional changes in *P. pectinata* in three notable genes: *Crk-l*, an integrator of multiple signaling pathways; Annexin A1 (*Anxa1*), a modulator of FGF ligands and RAS, and FGF1 intracellular binding protein (*Fibp*), an EGF/MAPK modulator (Fig. 2D-H)^19,20^. CRK-L had decreased hydrophobicity in the Src Homology 2 (SH2) domain where tyrosine phospho-proteins like growth factor receptors bind and decreased hydrophobicity in its SH3 domain that associates with RAC1 or RAS^21^ (Fig. 2D). ANXA1 had changes in all four Annexin repeat domains, which upon Ca^2+^ binding display phosopho-sites (Fig. 2E). Annexins activate MAPK signaling either through growth factors, or they can be directly phosphorylated by RTKs^22^. FIBP, which binds FGF1, showed fluctuations in both hydrophobicity and polarity in the annotated FIBP domain, though little is known about the mechanisms of action of this protein^23^ (Fig. 2F). Selection in genes such as these, which are potentially involved in body patterning and growth factor signaling suggest a role related to rostral development. A prime example is *Crk-l*, which has been shown to cause craniofacial defects in mice through disruption of neural crest development or through the retinoic acid/*Tbx1* regulatory network^24–26^.

Another major function of Cluster 1 genes is microtubule biology with two genes being significant (*Haus2* and *Tubgcp4*). Shared evolutionary trajectory with developmental regulators suggests a similar functionality in sawfish biology, such as through cilia-mediated signaling. One example is *Ccdc103*, which is necessary for ciliogenesis.

CCDC103 had numerous changes in polarity and hydrophobicity in its Rpap3 domain, which binds the axonemes of cilia (Figs. 2G). This is necessary for outer dynein arm attachment, and thus changes in this domain suggest functional differences which do not appear to be a result of gene duplication and subsequent divergence^27^. Lastly, a homeobox transcription factor involved in segmentation and body patterning, *HoxA5*, was found. The protein sequence had changes in polarity and hydrophobicity near the conserved site (residues 183–188) and the beginning of the DNA-binding homeobox domain, despite being truncated relative to *L. erinacea*. These changes suggest modified interactions between HOXA5 and its cofactors and/or targets (Fig. 2H).

To support whether changes in PSGs were specific to *P. pectinata* or shared with other sawfishes, sequencing of genomic DNA in functional domains was performed using samples from three other *Pristis* sawfishes: dwarf sawfish (*P. clavata*), green sawfish (*P. zijsron*), and largetooth sawfish (*P. pristis*). *Ccdc103* and *HoxA5* were chosen as they demonstrated clear chemical differences between *P. pectinata* and *L. erinacea* but were conserved enough in flanking regions of functional domains for amplification. Sequence alignments revealed that the ratios of non-synonymous to synonymous nucleotide substitutions in both genes were lower among sawfishes than when comparing sawfishes to *L. erinacea*. At *HoxA5* conserved sites, all five substitutions were shared among sawfishes but were non-synonymous with *L. erinacea*, with amino acid property changes at two sites (Fig. 2L, Sup. Fig. 3). A similar pattern was seen in *Ccdc103* among sawfishes. In the Parp3 domain of *Ccdc103*, 18/21 changes were non-synonymous between sawfishes compared to *L. erinacea*, and eight had an amino property change (Fig. 2J, Sup. Fig. 4). Fewer changes were seen among sawfishes; 10/13 were non-synonymous, and seven of these included chemical changes. Given the functions of PSGs and that the ratios of non-synonymous substitutions were lower among *Pristis* sawfishes compared to *L. erinacea*, this supports that these changes could be related to themes found in Cluster 1 genes. As conservation of the biochemical changes occurred at the base of the *Pristis* branch and conserved in all species this leads to further credence for a role in sawfish-specific morphological development.

**Figure 3.**
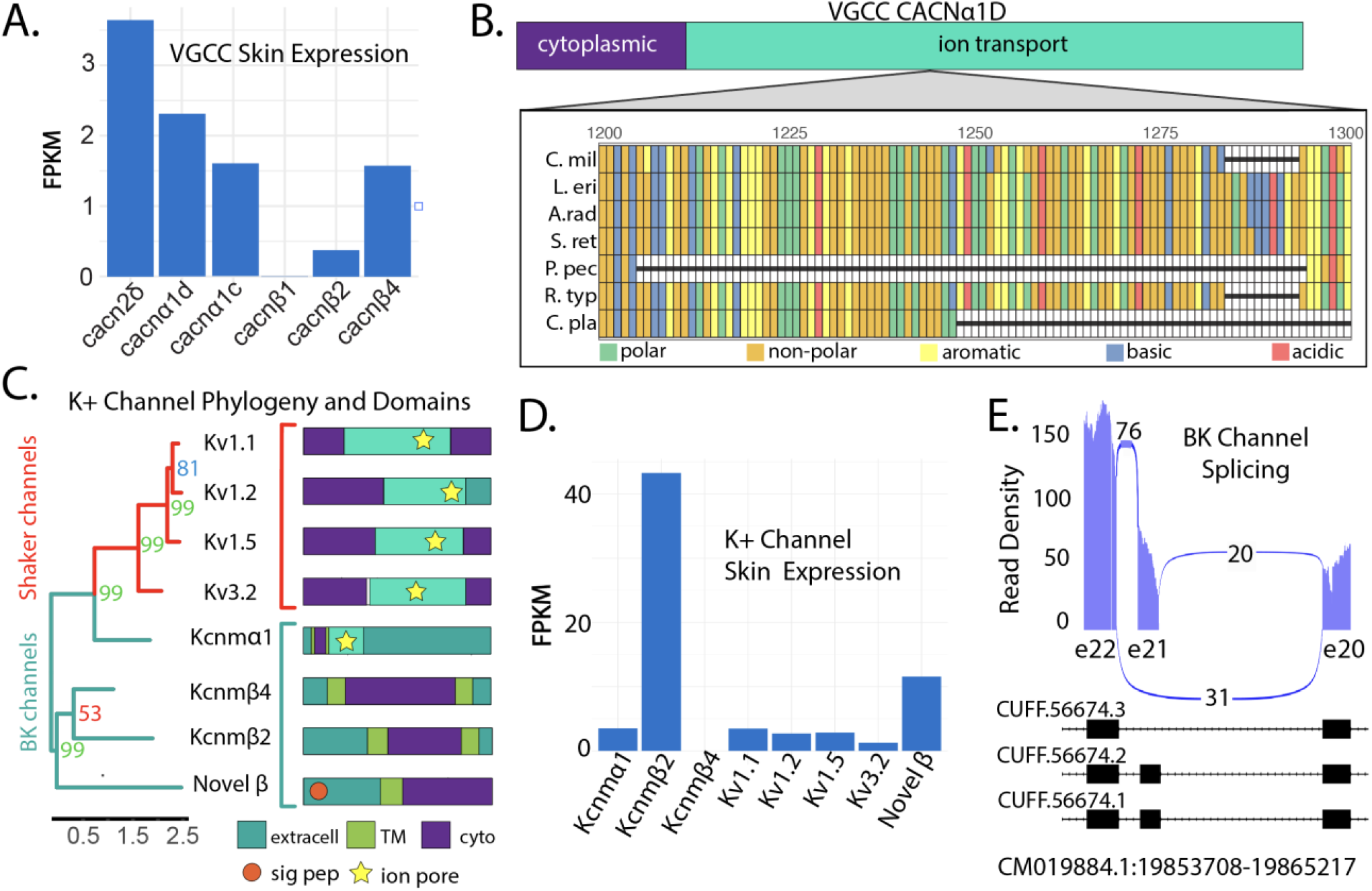
Smalltooth sawfish, *Pristis pectinata*, electroreception machinery. **A)** FPKM normalized expression of voltage gated calcium channel α and β subunits in rostral skin. **B)** Multiple sequence alignment of *Cacn*α*1D* depicting missing 91-residue segment in *P. pectinata* ion channel domain relative to other elasmobranchs. Species included: *Callorhinchus milii* (C. mil), *Leucoraja erinacea* (L. eri), *Amblyraja radiata* (A. rad), *Scyliorhinus retifer* (S. ret), *P. pectinata* (P. pec), *Rhinchodon typus* (R. typ), and *Chiloscyllium plagiosum* (C. pla). Alignments are colored by amino acid chemistry. **C)** Phylogenetic construction of potassium channels and subunits with bootstrap values noted (left). Sequences were aligned with ClustalW and tree was constructed by RAxML with 1000 bootstraps. Shaker channel sequences are highlighted in red, while BK channels are highlighted with light blue. Annotated protein domains of each sequence obtained using Interpro Scan (Right). EC is extracellular, TM, transmembrane, cyto is cytoplasm, and pore is ion channel pore. **D)** Normalized expression of potassium channels potentially involved electrosensing in rostral skin. Kcnm-transcripts correspond to BK channel subunits, Kv are voltage-gated Shaker channels. Novel β refers to the uncharacterized BK subunit found in transcriptome data. **E)** GGSashimi plot of alternatively spliced BK channel transcripts in all *P. pectinata* tissues. Transcriptome expression confirms that the BK channel transcript with highest expression in the rostral skin retains exon 21 (exon 29 in *L. erinacea*).

## Characterization of electrosensory genes

In sawfishes, the rostrum provides an exaggerated platform for placement of electrosensing organs called ampullae of Lorenzini. Initial sensing occurs through voltage-gated calcium channels (VGCC) produced from *Cacnα1D* in both rays and sharks^28^. Following influx of Ca2+ ions, K+ efflux occurs leading to membrane depolarization and neurotransmission. Different K+ channels participate in rays (BK-channels, BK-α) and sharks (Shaker-type, Kv). Using the *P. pectinata* transcriptome collection described above, orthologs of all channels involved were identified to understand mediators of electrosensing in this species.

First, all VGCC subunit transcriptomes were assessed for expression in rostral skin (Fig. 3A). Six VGCC-related genes were expressed, including a *Cacnα1D* ortholog. Others were channel accessory subunits (*Cacnα2δ, Cacnb2*, and *Cacnb4*), or a *Cacnα1C*-type VGCC, which have not been previously implicated in electrosensing chondrichthyans. A *Cacnb2* ortholog was found in the transcriptome but was not expressed in rostral skin. Thus, as expected, the *Cacnα1D* ortholog is likely the primary mediator of electrosensing. However, *P. pectinata Cacnα1D* is missing a 91-residue segment that would interact with intracellular effectors which is present in other rays (Fig. 3B)^28^. A portion of this region is also deleted in other chondrichthyans such as *C. mili* and the whale shark, *Rhincodon typus*, indicating that it is a site of functional novelty. Together, these results suggest that substantial changes occur during the initial step in the ability of *P. pectinata* to detect the bioelectric signal of prey.

Next, sequences of eight orthologs of Ca^2+^ responsive K^+^ channel were identified and compared (Fig. 3C). Clustering showed both major groups, BK (α and b) and Shaker types, were present in the gene set. A BK-α ortholog with clear pore motif was found (*Kcnmα1*) along with three accessory BK-β subunits, one of which was designated as novel as it could not be definitively paired with an ortholog. The four Shaker-type channels also exhibited the requisite pore domain residues. Two of these channels have been observed in *L. erinacea*, but they have only been demonstrated to work in conjunction with *Cacnα1D* orthologs in ampullae of the chain catshark, *S. retifer*, though they have also been found to be enriched in ampullae of the American paddlefish, *Polyodon spathula* ^29,30 31^.

All eight channel components are expressed in rostral skin, except for the *P. pectinata* β-4 ortholog (Kcnmb4) (Fig. 3D). Absence of this inhibitory subunit may contribute to the heightened electrosensing seen in sawfishes. Further differences were seen in expression levels of β-2 subunits. In *L. erinacea*, expression of β-subunits in electrosensory cells is 1000-to 10000-fold lower than α-subunit expression, yet in *P. pectinata* tissue, a BK β-2 is expressed nearly 10-fold to BK-α, and the novel subunit nearly 2-fold^29^. Together, this suggests *P. pectinata* has altered BK-channel physiology. Another unusual observation was expression of all four shaker-type channels, which differs from *L. erinacea* where only two, Kv1.1 and Kv1.5, are found^29^.

Other aspects of electrosensing machinery in *P. pectinata* showed potentially divergent biology. Several inserts and deletions were found in *Kcnmα1* that may alter its function (Sup. Fig. 5). The ion pore-containing BK α-subunits can differentially associate with β-subunits to modulate membrane repolarization to alter activation rates or affect calcium sensitivity, drawing into question the role of the novel β-subunit in the overall performance of the system^32–34^. Another difference found in *Kcnmα1* was the inclusion of exon 21 (known as exon 29 in *L. erinacea*) (Fig. 3E). Alternative splicing at exon 21 has been identified in BK channel ampullary transcripts of *L. erinacea* and in auditory hair cells of chick cochlea, though the effect on electrosensing is not known ^35^. The inclusion of this exon appears to be variable among elasmobranchs, being present in *P. pectinata*, thorny skate, *Amblyraja radiata*, and *L. erinacea* adult ampullae, but absent in *S. retifer*, whitespotted bamboo shark, *Chiloscyllium plagiosum*, and *L. erinacea* embryonic sequence (Sup. Fig. 5C).

## Implications of genetic changes in sawfishes

Here, fundamental changes in developmental genes were found in *P. pectinata*, which may influence morphogenesis of the rostrum. Elongated rostral structures have evolved in chondrichthyans at least five times^36^. Sawfishes (Batoidea: Pristidae) and sawsharks (Selachii: Pristiophoridae) are the only extant families that possess toothed rostra, but similar structures are also found in the fossil record, including two species of chimaeras, Squaloraja *polyspondyla* and *Acanthorhina jaekel* (Holocephali), Sclerorhynchoidei (Rajiformes), and *Bandringa rayi*. (Elasmobranchii)^36–38^. Convergent evolution of elongated rostral structures suggests an evolutionary advantage in prey detection, likely through heightened bioelectric sensing, which may provide sensory information at night, at depth, or in murky shallow waters^39^. The cohort of genes uniquely under selection in *P. pectinata* may provide a framework for the genetic changes that underpin the amorphogenetic origins of saw-like rostral structures. Primarily, this appears to be through reshaping the signaling environment, as was observed in this study, with changes in FGF/MAPK mediators. Simultaneously, positive selection was also seen in *HoxA5*, a transcription factor that establishes regions along the dorsoventral axis. However, in vertebrate development, cluster 5 Hox genes are typically expressed in the somites which eventually become the upper thorax and organs, and thus changes in this gene in *P. pectinata* may be related to differences in pectoral or synarcual morphology, and not related to the rostrum.

A consequence of potentially altered FGF/MAPK signaling in *P. pectinata* is divergent cellular response to environmental stressors such as changes in salinity, temperature, and dissolved oxygen. In Florida, juveniles have an affinity for warm (>30°C), brackish waters (18–30) with high dissolved oxygen levels (>6 mg/L).^40^ Cold water temperatures (<8–12°C) are known to alter habitat use and cause mortality^40,41^, while the physiological impacts of hypoxia and low salinities are unknown. As *P. pectinata* begins to recover, populations should re-establish in historical habitats. While there is evidence that such re-expansions have begun in the northern Gulf of Mexico and in the southern Indian River Lagoon, Florida^42^ (G. R. Poulakis unpublished data), water quality issues and fluctuating environmental conditions, caused in part by freshwater diversions that modify the hydrology of estuaries and cause algal blooms and hypoxic conditions, may pose a risk to sawfish health and viability^42^.

The genes involved in this environmental signaling can be affected by marine pollutants including polycyclic aromatic hydrocarbons (PAHs), heavy metals, bisphenol-A’s (BPAs), brominated flame retardants (BFRs), and polychlorinated benzodioxins (PCBs). For example, *Crk-l* expression is affected by BPAs, BFRs, and PCBs, and *Anxa1* is upregulated in response to PAH exposure in sea turtles as a response to increased production of reactive oxygen species^43^. Cadmium, PCBs, and other persistent organic pollutants can bioaccumulate in elasmobranch tissues, and may disrupt critical processes such as metabolism, immune function, and reproduction, though pollutant levels in sawfish tissues have not been reported^44–46^. Future research should identify contaminant levels in sawfish tissues and document sources of these pollutants in habitats where *P. pectinata* reside and may be re-establishing. Characterizing direct physiological effects of pollutants in sawfishes via *ex situ* experiments is not feasible; however, a potential circumvention of this issue could be to identify biomarkers in *P. pectinata* which are correlated with toxicological risks^47^.

A top conservation priority for *P. pectinata*, and all sawfishes, is to reduce injuries and mortalities in fisheries, especially trawl fisheries. Understanding the basis of sawfish electrosensing at a molecular level may support the development of more effective deterrent technologies that exploit this sensing modality. In future studies, molecular-level responses to stimuli, such as electric fields, should be studied to refine optimal experimental conditions, and used in parallel with aquarium trials to elicit avoidance responses^12^. Analysis of electrosensory genes revealed numerous differences in functional domains and expression of channels and subunits which were previously not implicated in electrosensing in chondrichthyans. As basal batoids, sawfish may possess unique electrosensing biology that leverages both types of K+ channels found in sharks and other rays. Considering ray-type electrosensing alone, elevated expression of multiple previously uncharacterized subunits suggests *P. pectinata* may have substantially altered BK-channel physiology. Altogether, these results suggest unique physiology of ion channels and subunits in *P. pectinata*, and their identification could allow tailored bycatch-reduction technology. Developing this technology for trawls is a priority as shrimp trawls have been identified as still having the highest bycatch risk for sawfishes in the ‘stronghold’ nations of the U.S. and Australia^9^.

In addition to providing an annotation of the publicly available genome, this study highlights the potential value of genomic approaches to conservation efforts, particularly through the identification and characterization of electrosensory genes and subunits and genes that could be used to reduce bycatch. Global sawfish recovery requires aggressive conservation planning that uses novel methodological approaches to support modern, high-tech solutions. Results from this work suggest that the impacts of pollutants need to be more deeply investigated and that physiological responses may differ between sawfishes and existing model organisms. These data will also support the ability to build experimental systems to test channel behavior in a controlled *in vitro* setting that can be used to assess and quantify the performance of sawfish electrosensing to facilitate development of behavior-modifying technology. Together, these data and insights provide the foundation to support key future research, with the goal of supporting global recovery of imperiled sawfishes.

## Supplementary methods

### Sample collection

RNAs were collected from the brain, liver, kidneys, ovary, and skin tissues of a juvenile female *P. pectinata*. Tissues were preserved in RNAlater and stored at −80°C until use. Samples from each tissue were homogenized and extracted with the TRIzol method. Concentrations and purities of RNA extracts were measured by nanodrop and bioanalyzer 2100 (Sup. Tbl. 1).

### Transcriptome sequencing and assembly

The transcriptome was assembled using Trinity on the Magnolia High Performance Computing (HPC) cluster^48^. Blast2GO was used to assign gene ontology terms to transcripts, and WeGO was used for comparison between *P. pectinata* and *L. erinacea*^49,50^. Completeness was assessed with BUSCO in transcriptome mode against the most recent vertebrate lineage, odb10. To annotate the genome with coding sequences and to capture a more complete gene set for positive selection analysis, the published *P. pectinata* genome from an adult female (GCA_009764475.2) was used in addition to the assembled transcriptome.

### Genome annotation for positive selection analysis geneset

Gene content exclusive to the transcriptome was isolated using STAR aligner with default parameters to align reads from all tissue samples to the genome^51^. Coverage over the genome was assessed with BEDtools and reads with low coverage (< 5 reads) were then aligned to the assembled transcriptome using Hisat2 with the –dta-cufflinks option enabled^52,53^ Cufflinks was used to map RNA reads to coordinates in the genome, which were then intersected, excluding overlapping transcripts, with Augustus-predicted gene sequence coordinates for the genome using BEDTools Intersect^54^. All BAM and SAM file conversion and sorting was performed with SAMtools^55^. Unique gene sequences from the transcriptome and genome were concatenated into one dataset for further analysis and BUSCO annotation output was used to remove redundant sequences and assess completeness. To obtain transcript expression, each tissue library was individually mapped to the constructed transcriptome using Bowtie2, and SAMtools Idxstats was used to quantify mapped reads^56^.

### Positive selection in transcriptome

Open reading frames were predicted from all input transcriptomes using TransDecoder with default parameters^57^. All coding sequences were concatenated into one dataset and Orthofinder was executed with default parameters to cluster orthologous gene groups^16^ Sequences in each gene family were annotated using eggNOG with Diamond mode enabled^58^. Annotations were also used to split gene families into paralogous groups using custom scripts. For positive selection, only groups that retained at least one sequence from each taxon were retained for analysis. Transcripts were discarded if their length was more than 100 amino acids shorter or longer than the average length of the gene, and only genes with Pal2Nal alignments longer than 20 amino acids were kept, excluding trees with insufficient branch lengths for analysis. ABSREL, which identifies episodic selection in individual branches, was used to analyze each group of orthologous genes^17^. Mafft v7.475 was used for protein alignment, FastTree 2.1.10 for tree construction, and Pal2Nal v14 with the –nogap option to provide gap-free codon-based nucleotide alignments^59–61^. Output JSON files were parsed with custom scripts using the JsonLite package^62^. To examine potential effects of substitutions in genes of interest under selection, chemical properties were compared between protein sequences from *P. pectinata* and *L. erinacea* using Expasy ProtScale^63^. The Kyte & Doolittle (1982) scale was used for hydrophobicity and the Zimmerman scale for polarity. After manual gap correction, scale values at each residue for *P. pectinata* were subtracted from *L. erinacea* values and the change plotted by residue. Domains of each protein were obtained from Interpro Scan. Analysis of PSGs of interest using DAMBE found no significant substitution saturation in any species from any alignment, indicating that the species are not too diverged to obtain meaningful positive selection results^64^.

For PCA analysis, grand average values of hydropathicity (GRAVY) for each *P. pectinata* and *L. erinacea* protein were obtained from Expasy ProtParam. If there were two sequences in an orthogroup for *L. erinacea*, the closest aligning sequence was selected for each *P. pectinata* gene from a multiple sequence alignment, and the difference was taken between the values. Expression of homologous genes under selection in *P. pectinata* was compared to similar tissues from *S. retifer* by aligning publicly available single-end RNAseq libraries from brain, liver, kidney, ovary, and skin to the longest *S. retifer* homolog sequence (Sup. Tbl. 4). Alignment was performed with Hisat2 and quantified with SAMtools Idxstats^52,55^. Alignment statistics for each tissue type can be found in Sup. Table 5. Expression values for each gene were normalized to TPM and subtracted from *P. pectinata* values. Principal component analysis was performed using the factoextra R package and visualized with ggpubr^65^ (Supplementary File 1).

### Conservation of changes in sawfishes

To determine whether signals of positive selection in genes of interest for *P. pectinata* were also present in other sawfish species, nucleotide changes in the three other *Pristis* sawfishes were examined for HoxA5 and Ccdc103. Primers were designed to amplify divergent homologous functional domains between *P. pectinata* and *L. erinacea*. Primer sequences used were: *HoxA5* forward: 5’GACTTATGTGCAGTTTTCGCATCCA 3’; *HoxA5* reverse: 3’AACTACCTCCTCAAATTC 5’; *Ccdc103* forward: 5’CTGCTGCTCAGGAAATCCAC 3’; *Ccdc103* reverse: 3’AGCGGAGTTTAGCCGTGACTG 5’.

DNA samples from *P. clavata, P. zijsron*, and *P. pristis* were mixed with Phire Hot Start II DNA polymerase (ThermoFisher), dNTPs, deionized water, and primers for either *Ccdc103* or *HoxA5*, followed by PCR amplification using a Mastercycler Pro and electrophoresis apparatus. The reactions were amplified for 35 cycles at 98°C for 30s/5s, 53°C/58.5°C for 15s, and 72°C/72°C for 1min/1 min for *HoxA5* and *Ccdc103* respectively. The agarose gel bands were purified with a GeneJET Plasmid MiniPrep (ThermoFisher) and sent to Eurofins Genomics for sequencing. ApE (RRID:SCR_014266) was used to translate DNA sequences into the appropriate reading frame, and Clustal Omega was used to align sequences to *P. pectinata* and *L. erinacea*^66^. BLAT from the UCSC genome browser and the *P. pectinata* genome were used to verify amplification of the correct target sequence. Alignments were manually examined between species to identify conserved non-synonymous changes among sawfishes that were not found in *L. erinacea*.

### Electrosensory genes

Full coding sequences for electrosensing genes were downloaded from NCBI and used for manual positive selection analysis. Analysis included the VGCC (*Cacnα1D*) and several β-subunits, potassium-activated (BK) channel α subunit, and several Shaker (Kv) channels which have been implicated as major ion channels involved in electroreception in sharks and rays^28^ Transcripts for *P. pectinata* were identified with BLASTP using *L. erinacea* sequences from NCBI (acc. AJP74816.1, KY355736.1) as query and a protein database was constructed from the transcriptome by Hmmer2GO^67,68^. BK*β* and Shaker-type channels were identified using BLAST and confirmed using Interpro Scan^69^. Multiple sequence alignments were performed using ClustalW with default parameters and 1000 bootstraps and visualized using the GGMsa package in R^70^. The phylogenetic tree was visualized with the GGTree package^71^. GenBank accessions for the elasmobranch sequences most similar to the uncharacterized BK*β* subunit were: XP_041.47886.1, XP_032892500.1, GCB66272.1, XP_038668782.1, and XP_043564107.1 (accessed 9/26/2022).

## Data availability statement

Raw sequence data used to assemble the transcriptome can be found under NCBI accession PRJNA864825.

## Author contributions

NP, KF, and AF administered the project. GN contributed the high coverage reference genome assembly. AF, TJ, and NP designed and executed experiments. TJ and AF analyzed and interpreted data. CVE performed PCRs for *Ccdc103* and *HoxA5*. GP acquired and maintained the endangered species collection permit and collected the tissue samples from *Pristis pectinata*. TJ, AF, NP, GP, and GN wrote the manuscript and/or contributed to final edits.

## Acknowledgments

Funding for this project was provided by NOAA Awards NA16NMF4720062 (field work) and NA18NMF0080237 (laboratory processing), start-up funds from the University of Florida to GN for the long read sequencing and assembly of the reference genome. The authors acknowledge HPC at The University of Southern Mississippi supported by the National Science Foundation under the Major Research Instrumentation (MRI) program via Grant #ACI 1626217. Thanks to David Morgan, Jeff Whitty, and the Western Australia Department of Fisheries for providing tissue samples of *P. clavata, P. pristis, and P. zijsron*.

---

**Supplementary Figure 1.**
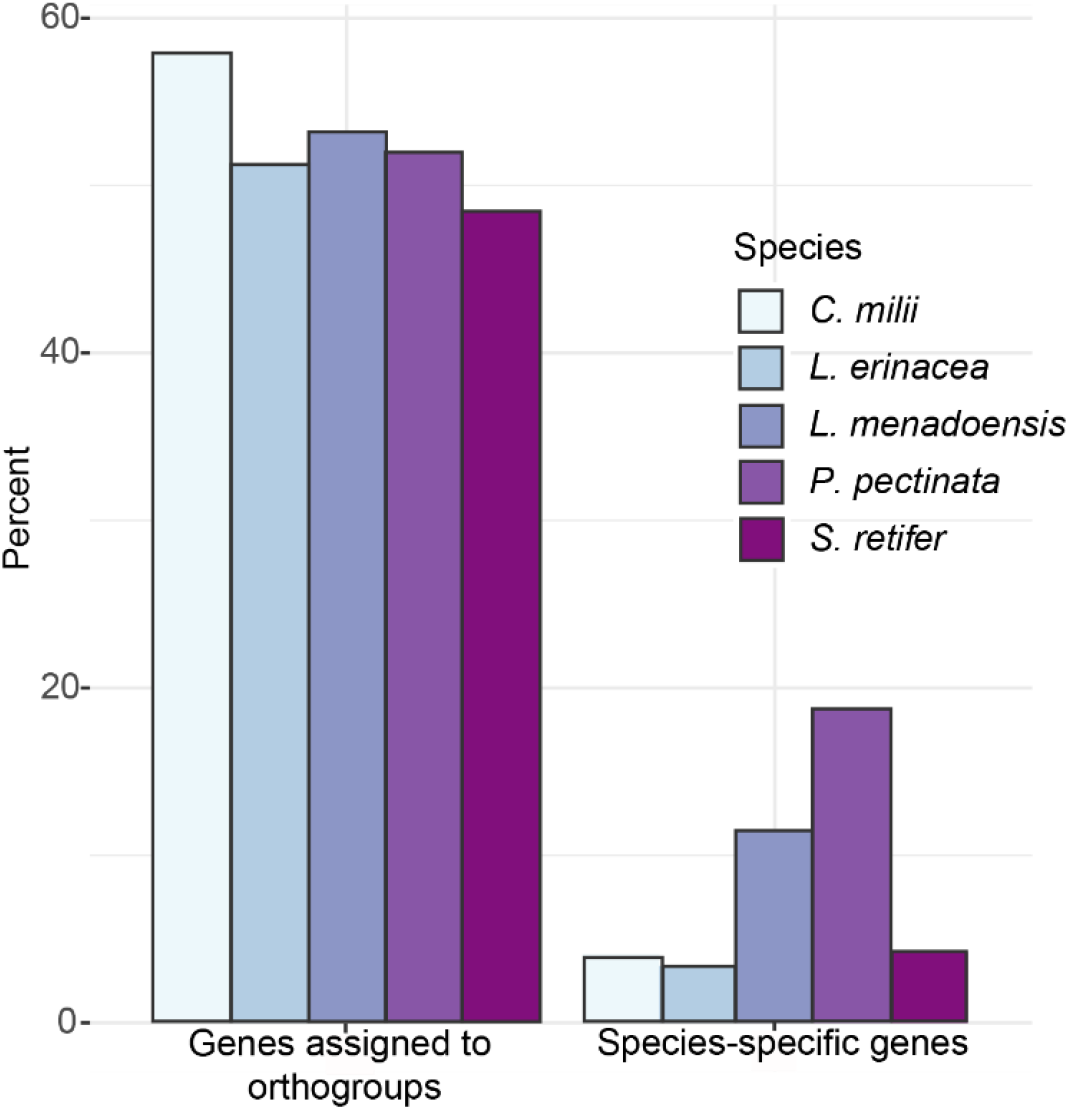
Left: Percentage of genes assigned to orthogroups by OrthoFinder. Right: Percentage of genes unique to each species. Species included: chain catshark, *Scyliorhinus retifer*, Indonesian coelacanth, *Latimeria menadoensis*, Australian ghostshark, *Callorhinchus milii*, smalltooth sawfish, *Pristis pectinata*, and little skate, *Leucoraja erinacea*.

**Supplementary Figure 2.**
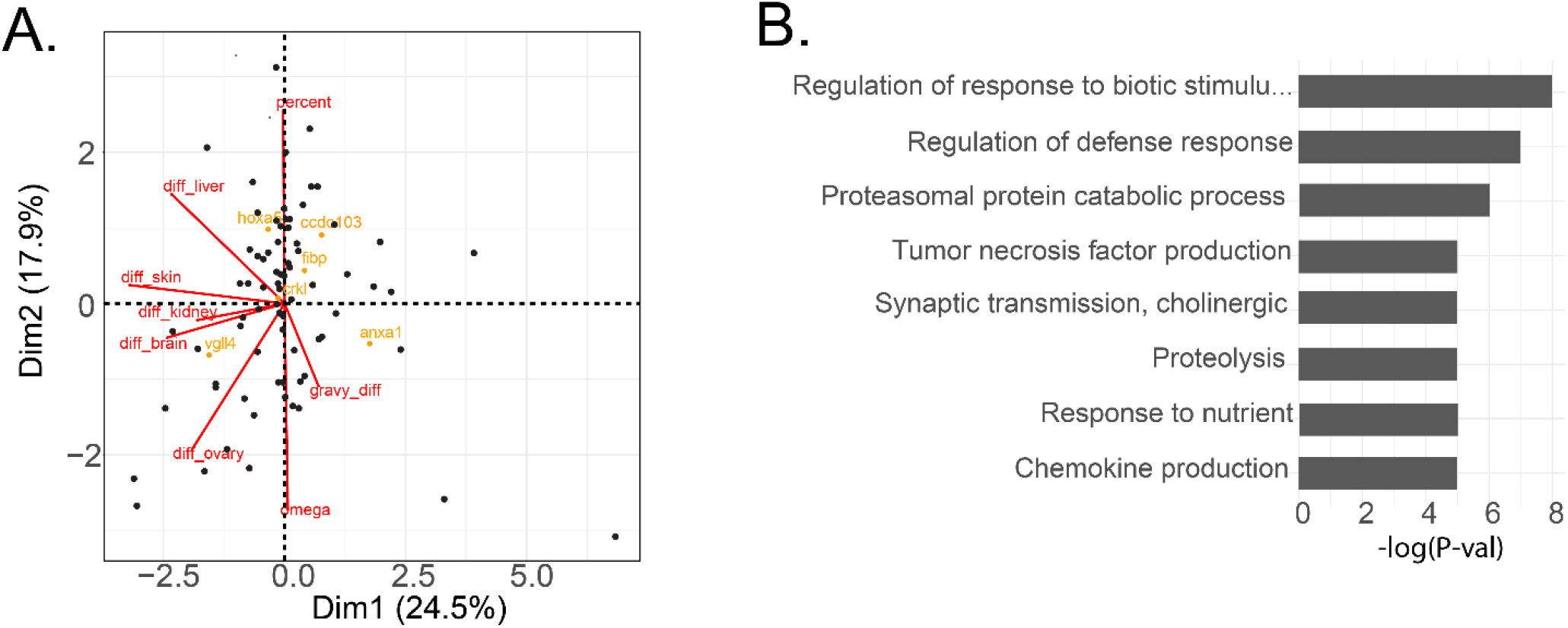
Smalltooth sawfish, *Pristis pectinata*, genes under selection. **A)** PCA of genes under selection using omega value, percent of sites, change in GRAVY value relative to little skate, *Leucoraja erinacea*, and change in expression relative to chain catshark, *Scyliorhinus retifer*. Genes of interest from Cluster 1 are shown in yellow, variables are shown in red. **B)** Top 8 most enriched gene ontology terms related to biological processes from genes grouped into Cluster 2 by PCA analysis plotted by -log(p-value).

**Supplementary Figure 3.**
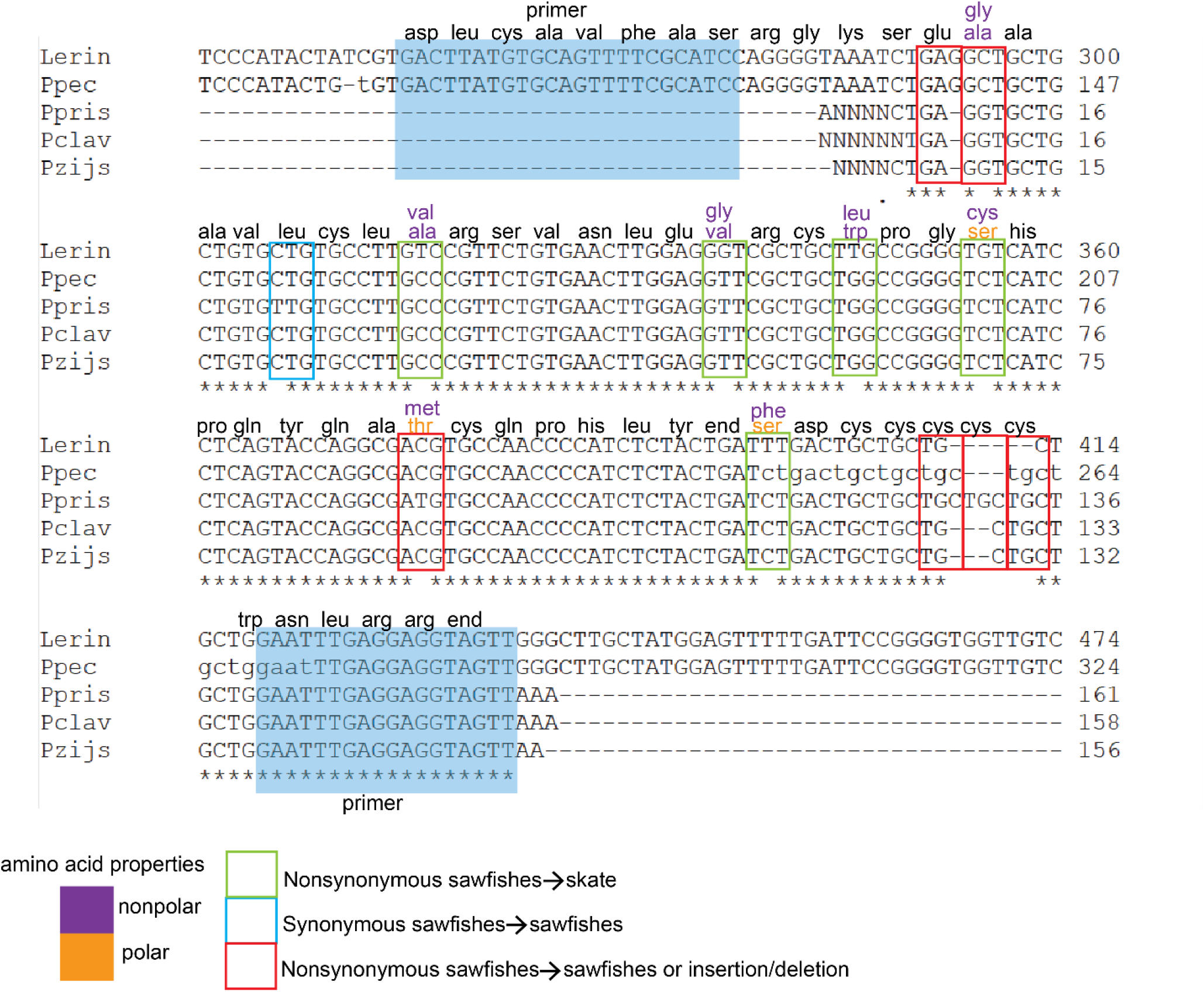
Annotation of substitutions in *HoxA5* between dwarf, green, largetooth, and smalltooth sawfishes (*Pristis clavata, P. zijsron, P. pristis, P. pectinata*, respectively) and little skate, *Leucoraja erinacea*. Changes in amino acids are noted above aligned nucleotide sequences. Primer sequences are shaded in blue.

**Supplementary Figure 4.**
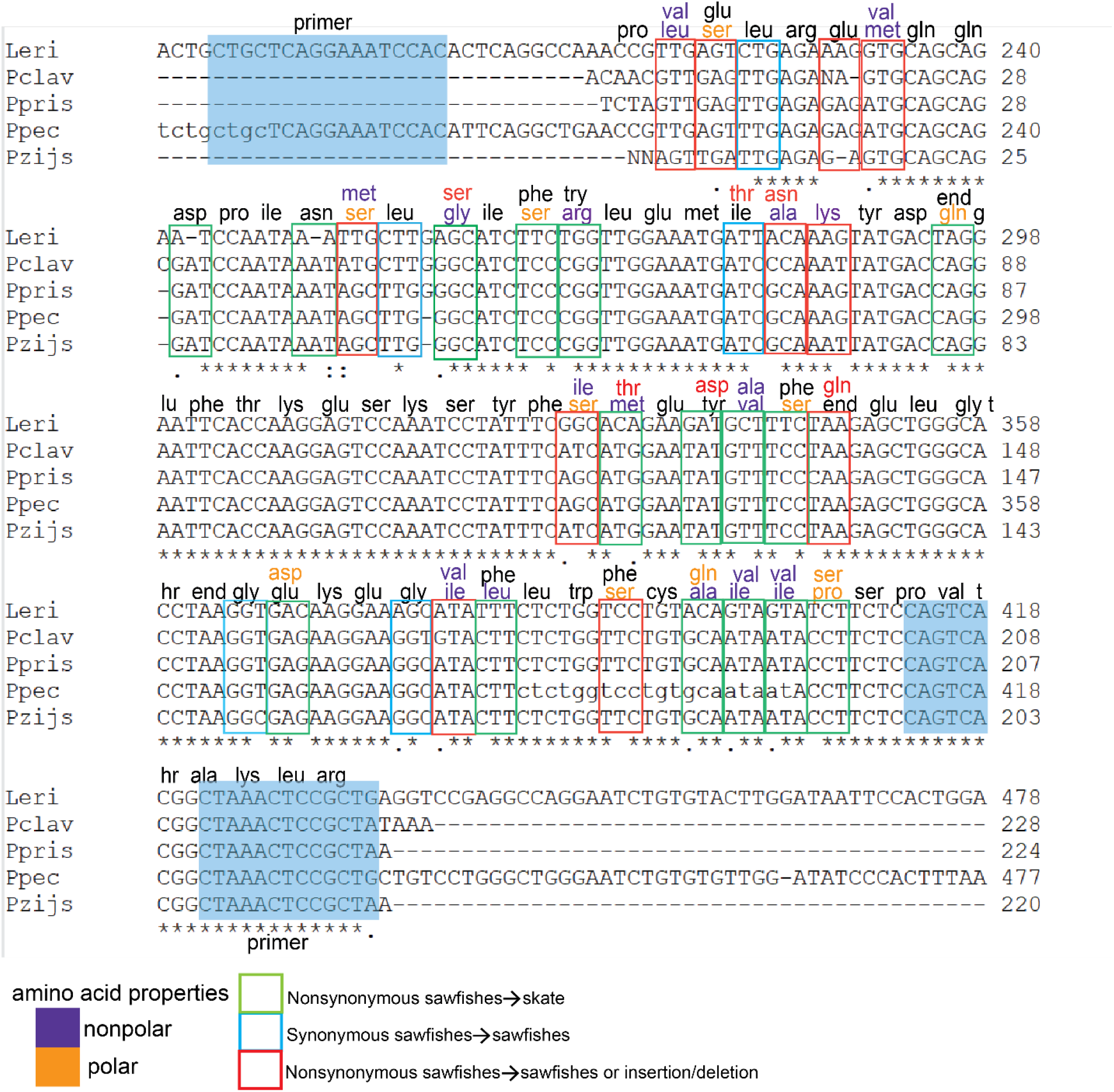
Annotation of substitutions in *Ccdc103* between dwarf, green, largetooth, and smalltooth sawfishes (*Pristis clavata, P. zijsron, P. pristis, P. pectinata*, respectively) and little skate, *Leucoraja erinacea*. Changes in amino acids are noted above aligned nucleotide sequences. Primer sequences are shaded in blue.

**Supplementary Figure 5.**
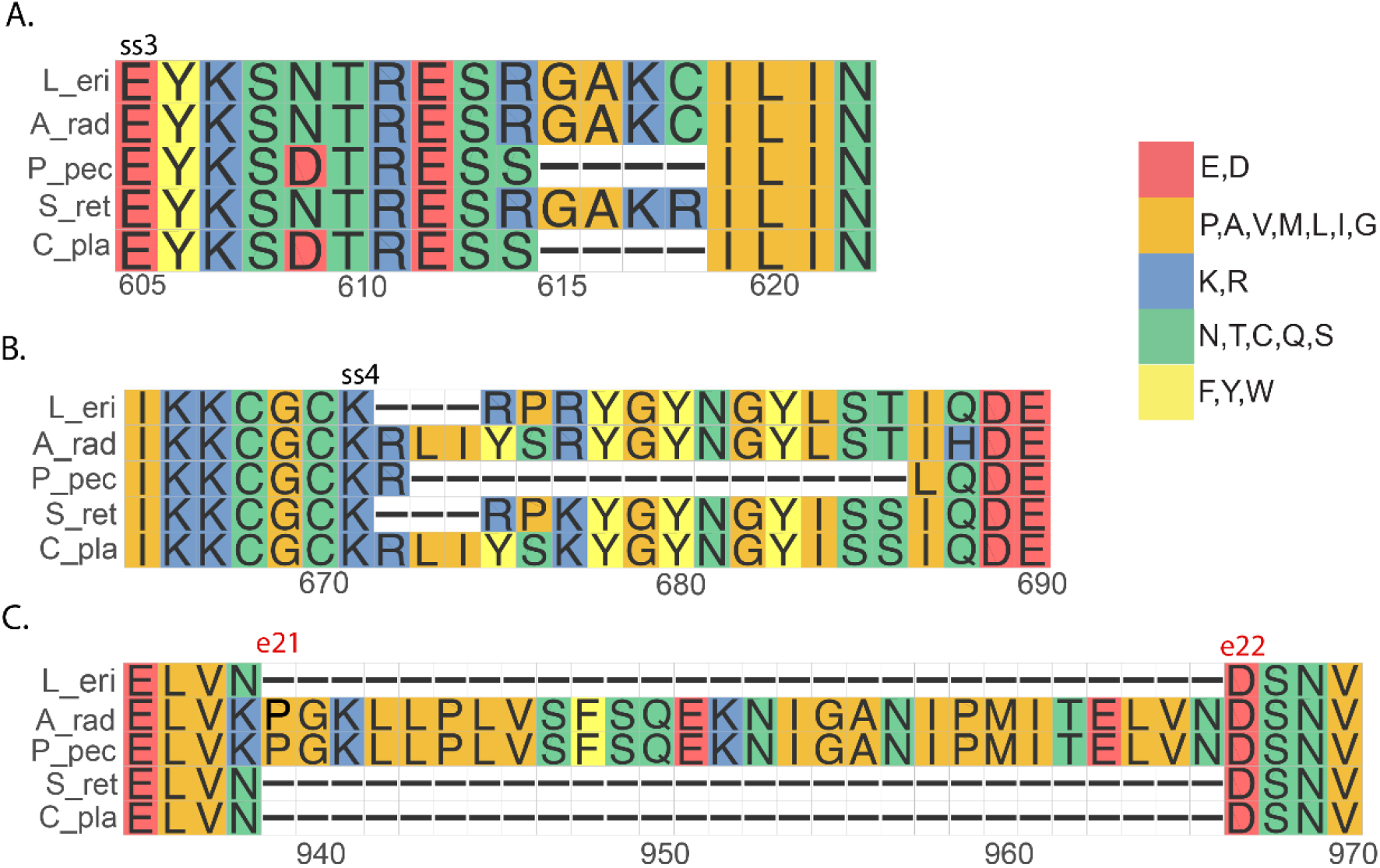
Insertions, deletions, and splice sites identified in *Pristis pectinata* (P_pec) BK alpha K^+^ channel relative to *Leuroraja erinacea* (L_eri), *Amblyraja radiata* (A_rad), *Scyliorhinus retifer* (S_ret) and *Chiloscyllium plagiosum* (C_pla). Ss = splice site. e = exon. Plot colors indicate amino acid chemical properties.

**Supplementary Table 1.** Concentrations and purities of smalltooth sawfish, *Pristis pectinata*, RNA samples by tissue obtained by nanodrop and Bioanalyzer. For RNA, chemical purity is indicated by A260/A280 of 2.0. RNA integrity number (RIN) ranges from 1 to 10, where 10 is intact and 1 is degraded.

| Tissue | Concentration<br>(ng/ul) | Purity<br>(A260/A280) | RIN |
| --- | --- | --- | --- |
| Brain | 174.7 | 1.83 | 6.7 |
| Liver | 962.4 | 2.01 | 6.9 |
| Ovary | 1660.7 | 2.03 | 7.0 |
| Kidney | 1150.0 | 2.04 | 7.3 |
| Skin | 273.9 | 1.98 | 7.5 |

**Supplementary Table 2.** Number of transcripts, NCBI accession numbers, and percentage of complete (C) and fragmented (F) orthologs via BUSCO analysis of transcriptomes for all taxa used in positive selection analyses.

|  | Smalltooth<br>sawfish,<br>*Pristis*<br>*pectinata* | Chain<br>catshark<br>*Scyliorhinus*<br>*retifer* | Little skate,<br>*Leucoraja*<br>*erinacea* | Indonesian<br>coelacanth,<br>*Latimeria*<br>*menadoensis*, | Australian<br>ghostshark,<br>*Callorhinchus*<br>*mili* |
| --- | --- | --- | --- | --- | --- |
| No.<br>transcripts<br>in dataset | 175,569 | 107,231 | 103,996 | 66,138 | 92,334 |
| Accession | NA | GEO:<br>GSM643958 | GEO:<br>GSM643957 | GAPS01066138 | GEO:<br>GSM643959 |
| %BUSCO<br>complete | 89.47% C | 56.6% C | 60.4% C | 40.9% C | 47.3% C |
|  | 3.87% F | 22.1% F | 20.4% F | 18.9% F | 26.9% F |

**Supplementary Table 3.** Number of unique genes and total number of genes found under selection in aBSREL analysis by species.

| Species | Unique genes | Total genes |
| --- | --- | --- |
| Smalltooth sawfish, *Pristis pectinata* | 77 | 96 |
| Chain catshark, *Scyliorhinus retifer* | 26 | 38 |
| Australian ghostshark, *Callorhynchus milii* | 76 | 95 |
| Little skate, *Leucoraja erinacea* | 40 | 51 |
| Indonesian coelacanth, *Latimeria menadoensis* | 49 | 60 |

**Supplementary Table 4.** NCBI sample names, accession codes, and run identifiers for the chain catshark, *Scyliorhinus retifer*, expression data used in PCA analysis.

| Sample | Accession code | Run |
| --- | --- | --- |
| Adult 1 kidney | SAMD00098998 | DRR111789 |
| Adult 1 medulla | SAMD00098989 | DRR111780 |
| Adult 1 ovary | SAMD00098994 | DRR111785 |
| Adult 1 liver | SAMD00098997 | DRR111788 |
| Adult 3 skin | SAMD00099043 | DRR111834 |

**Supplementary Table 5.**
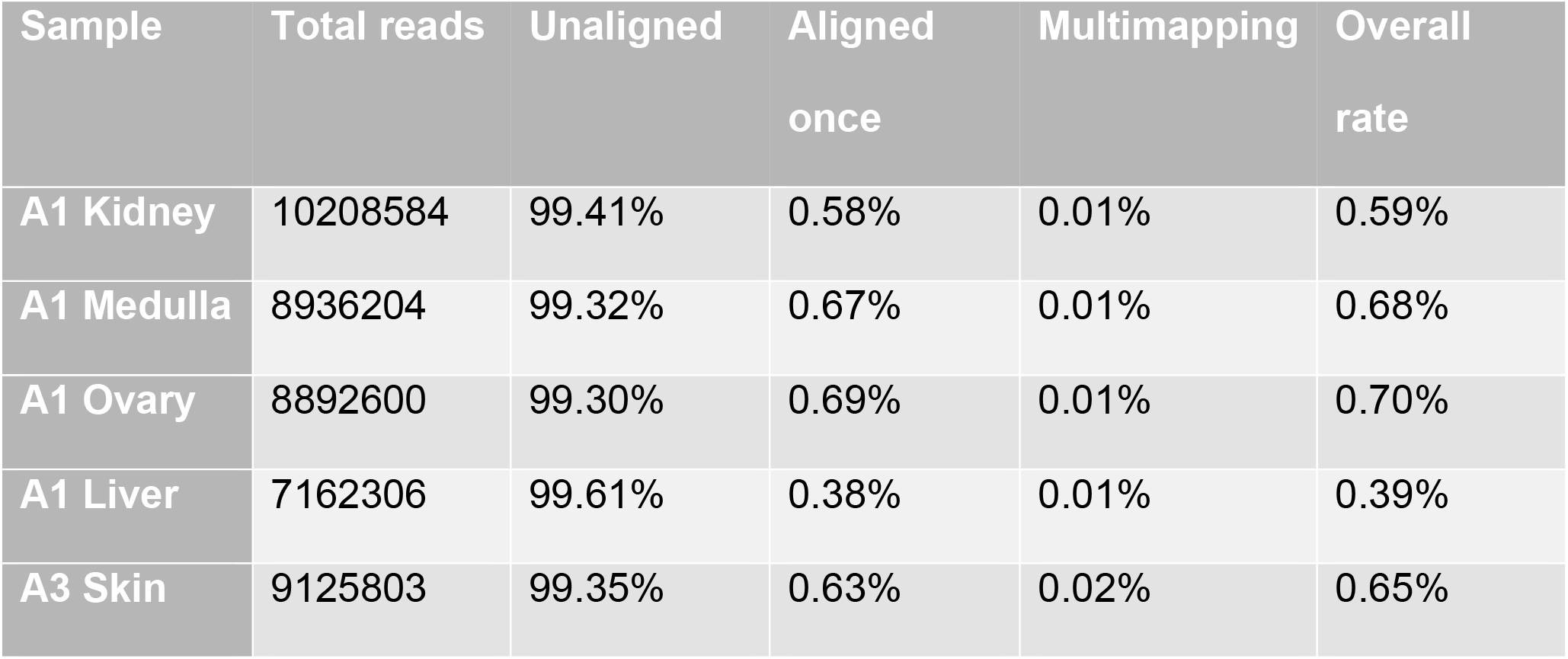
Number and percent of reads mapped to genes orthologous to the smalltooth sawfish, *Pristis pectinata*, PSGs per sample for the chain catshark, *Scyliorhinus retifer*, expression data used in PCA analysis.

**Supplementary Table 6.** BLAT results of Hox genes into the smalltooth sawfish, *Pristis pectinata*, genome browser with chromosome positions, starts, stops, highest scores, and query accession numbers. Only the highest scores were included for simplicity.

| Gene | Chrom | Start | Stop | Score | Query Acc. |
| --- | --- | --- | --- | --- | --- |
| hoxA1 | Chr5 | 107566999 | 107568411 | 902 | ACT65756.1 |
| hoxA2 | Chr5 | 107572840 | 107574441 | 1070 | ACT65755.1 |
| hoxA3 | Chr5 | 107579173 | 107581805 | 1056 | ACT65754.1 |
| hoxA4 | Chr5 | 107599253 | 107600338 | 706 | ACT65753.1 |
| hoxA5 | Chr5 | 107611902 | 107613201 | 706 | ACT65752.1 |
| hoxA6 | Chr5 | 107615218 | 107616103 | 680 | ACT65751.1 |
| hoxA7 | Chr5 | 107624042 | 107625644 | 611 | ACT65750.1 |
| hoxA9 | Chr5 | 107633224 | 107634655 | 610 | ACT65749.1 |
| hoxA10 | Chr5 | 107642407 | 107644791 | 686 | ACT65748.1 |
| hoxA11 | Chr5 | 107656057 | 107658036 | 818 | ACT65747.1 |
| hoxA13 | Chr5 | 107669970 | 107671270 | 860 | ACT65746.1 |
| hoxB1 | Chr25 | 3190815 | 3192527 | 718 | ASS31204.1 |
| hoxB2 | Chr25 | 3200448 | 3202103 | 728 | ASS31205.1 |
| hoxB3 | Chr25 | 3206913 | 3208894 | 1055 | ASS31206.1 |
| hoxB4 | Chr25 | 3228207 | 3229258 | 652 | ASS31207.1 |
| hoxB5 | Chr25 | 3242638 | 3244302 | 727 | ASS31208.1 |
| hoxB6 | Chr25 | 3246731 | 3248076 | 611 | ASS31209.1 |
| hoxB7 | Chr25 | 3254738 | 3256947 | 596 | XP_032890850.1 |
| hoxB8 | Chr25 | 3258510 | 3259634 | 611 | ASS31210.1 |
| hoxB9 | Chr25 | 3268829 | 3270311 | 493 | ASS31211.1 |
| hoxB10 | Chr25 | 3278434 | 3284468 | 488 | ASS31212.1 |
| hoxB13 | Chr25 | 3355373 | 3356805 | 793 | ASS31213.1 |
| hoxD1 | Chr1 | 47299641 | 47300923 | 482 | ASS31214.1 |
| hoxD2 | Chr1 | 47291801 | 47293231 | 808 | ASS31215.1 |
| hoxD3 | Chr1 | 47286156 | 47288710 | 929 | ASS31216.1 |
| hoxD4 | Chr1 | 47268047 | 47269186 | 623 | ASS31217.1 |
| hoxD5 | Chr1 | 47258508 | 47259603 | 566 | ASS31218.1 |
| hoxD8 | Chr1 | 47248080 | 47249022 | 665 | ASS31219.1 |
| hoxD9 | Chr1 | 47240275 | 47241612 | 590 | ASS31220.1 |
| hoxD10 | Chr1 | 47233908 | 47235988 | 715 | ASS31221.1 |
| hoxD11 | Chr1 | 47224113 | 47225493 | 592 | ASS31222.1 |
| hoxD12 | Chr1 | 47216240 | 47217755 | 701 | ASS31223.1 |
| hoxD13 | Chr1 | 47209657 | 47210874 | 773 | ASS31224.1 |

